# Using DNA from mothers and children to study parental investment in children’s educational attainment

**DOI:** 10.1101/489781

**Authors:** Jasmin Wertz, Terrie E. Moffitt, Jessica Agnew-Blais, Louise Arseneault, Daniel W. Belsky, David L. Corcoran, Renate Houts, Timothy Matthews, Joseph A. Prinz, Leah S. Richmond-Rakerd, Karen Sugden, Benjamin Williams, Avshalom Caspi

## Abstract

This study tested implications of new genetic discoveries for understanding the association between parental investment and children’s educational attainment. A novel design matched genetic data from 860 British mothers and their children with home-visit measures of parenting: the E-Risk Study. Three findings emerged. First, both mothers’ and children’s education-associated genetics, summarized in a genome-wide polygenic score, predicted parenting -- a gene-environment correlation. Second, accounting for genetic influences slightly reduced associations between parenting and children’s attainment -- indicating some genetic confounding. Third, mothers’ genetics influenced children’s attainment over and above genetic mother-to-child transmission, via cognitively-stimulating parenting -- an environmentally-mediated effect. Findings imply that, when interpreting parents’ effects on children, environmentalists must consider genetic transmission, but geneticists must also consider environmental transmission.

---

Parents devote a great deal of time and effort to ensuring their children’s educational success. They read to their children, buy educational toys, monitor their children’s schoolwork and enrol them in enriching classes and extracurricular activities. Such parental investment is partly motivated by the belief that what parents do is crucial for children’s educational success. However, this belief has not gone unchallenged. In popular books, pundits have questioned the importance of parental influence (Harris, 1998; Rowe, 1993) and lamented psychology’s focus on nurture over nature in shaping developmental outcomes (Pinker, 2002). In scientific journals, discussions continue about the relevance of parenting for children’s outcomes (Sherlock & Zietsch, 2018; Waldinger & Schulz, 2018). The debate about parental influences on children’s attainments has been fuelled by three lines of evidence from behavioral genetics research. First, genetic influences have been documented for all traits and behaviors, including children’s educational attainment (Asbury & Plomin, 2014; Polderman et al., 2015). Second, children’s genetics influence the parenting they receive. This is most apparent in research reporting greater similarity in received parenting among genetically identical versus non-identical twin children (Avinun & Knafo, 2014; Neiderhiser et al., 2004; Riemann, Kandler, & Bleidorn, 2012). Influences of children’s genetics on their received parenting come about because characteristics of children that are partly heritable elicit differences in parenting -- an ‘evocative’ gene-environment correlation (Plomin & Bergeman, 1991). Third, parents’ genetics influence the parenting they provide. This is most apparent in research documenting greater similarity in how identical versus non-identical adult twins parent their offspring (Klahr & Burt, 2014; Neiderhiser et al., 2004). Parents’ genetics influence parenting because parenting partly reflects personal characteristics that are themselves heritable. Parents’ genetic influence on their parenting creates an ‘active’ gene-environment correlation from the perspective of the parent (because parents’ genes will be correlated with the parenting they provide) and a ‘passive’ gene-environment correlation from the perspective of the child (because children will inherit genes that are correlated with the parenting to which they are exposed) (Plomin, DeFries, & Loehlin, 1977).

Gene-environment correlations in child development complicate the interpretation of socialization research (Scarr & McCartney, 1983). In particular, they raise the possibility that genetic influences confound associations between parenting and children’s educational attainment. This would be the case if genes that influence children’s educational attainment also affect the kind of parenting that is linked with educational success. Confounding could occur if parents’ education-associated genetics shape their parenting and are also passed on to their children in whom they influence children’s educational attainment. Confounding could also occur if children’s education-associated genetics influence both the parenting they receive and their educational attainment. In both of these scenarios, associations between parenting and children’s educational attainment may not reflect a causal effect of parenting on children. Instead, parenting may merely be a marker of children’s or parents’ education-associated genetic predisposition; in theory, it is possible that parenting lacks any environmental effects on children’s educational attainment of its own (Knafo & Jaffee, 2013; Moffitt, 2005). This possibility can be summarized as ‘genetic confounding’.

However, gene-environment correlations do not necessarily lead to confounding. Another possibility is that the portion of parenting that is genetically influenced still affects children’s educational attainment. This would be the case if parents’ genetics influenced how they parent, and parenting subsequently affects children’s educational attainment through environmental ways. Recent research has provided evidence supporting this possibility, showing that education-associated alleles of parents influence their children’s educational success, even if those alleles are not passed on from parent to child (Bates et al., 2018; Kong et al., 2018). This research ruled out genetic confounding by isolating the effects of parents’ education-associated alleles that were non-transmitted, i.e. not passed on to children. The findings suggest that parents’ genetics influence children’s educational outcomes via environments parents create. This possibility has been referred to as ‘genetic nurture’ (Kong et al., 2018). It implies that treating genetics as only a confounding influence on associations between parenting and child outcomes may leave behavioral scientists with an incomplete account of parenting effects on child development.

Here we used a novel design to test gene-environment correlations, genetic confounding and genetic nurture. Our design offers two innovative components. First, we computed genome-wide polygenic scores for <u>both</u> mothers and their children using genotype data that we collected from both generations. These families are participating in the Environmental Risk (E-Risk) Longitudinal Twin Study, a UK-based cohort study. Polygenic scores are derived from genome-wide association studies (GWAS; Visscher et al., 2017) and aggregate millions of genetic variants across the genome into a score that indicates part of a person’s genetic disposition to a particular trait or behavior (Dudbridge, 2013). Because the focus of this study is on parenting in relation to the outcome of children’s educational attainment, we calculated polygenic scores based on recent GWAS of educational attainment (Lee et al., 2018). The second design innovation was that we matched molecular-genetic data with extensive measures of mothers’ parenting that we collected in four successive family home visits during the first 12 years of children’s lives. Parenting measures were derived from multiple reporters: mothers, interview staff, and children themselves. We focused on aspects of parenting that have been shown to predict children’s educational attainment: cognitive stimulation; warm, sensitive parenting; low household chaos; and a safe, tidy home (Davis-Kean, 2005; Garrett-Peters, Mokrova, Vernon-Feagans, Willoughby, & Pan, 2016; Spera, 2005). We measured children’s educational attainment at age 18 years.

We used these data to test three hypotheses, as illustrated in Figure 1. First, we tested for the presence of gene-environment correlations. We did this by testing whether mothers’ education polygenic scores predicted the parenting they provided (Figure 1, Path a) and whether children’s polygenic scores predicted the parenting they received (Figure 1, Path b). Because mothers share genetics with their children (Figure 1, Path c), genetic associations with parenting could either reflect active gene-environment correlations between mothers’ genetics and parenting or evocative gene-environment correlations between children’s genetics and parenting. To disentangle active from evocative gene-environment correlation, we tested whether mothers’ polygenic scores predicted parenting after adjusting for children’s polygenic scores (indicating active gene-environment correlation) and whether whether children’s polygenic scores predicted parenting after adjusting for mothers’ polygenic scores (indicating evocative gene-environment correlation). A finding of positive gene-environment correlations would indicate that education-associated genetics shape the parenting mothers provide and children receive.

**Figure 1.**
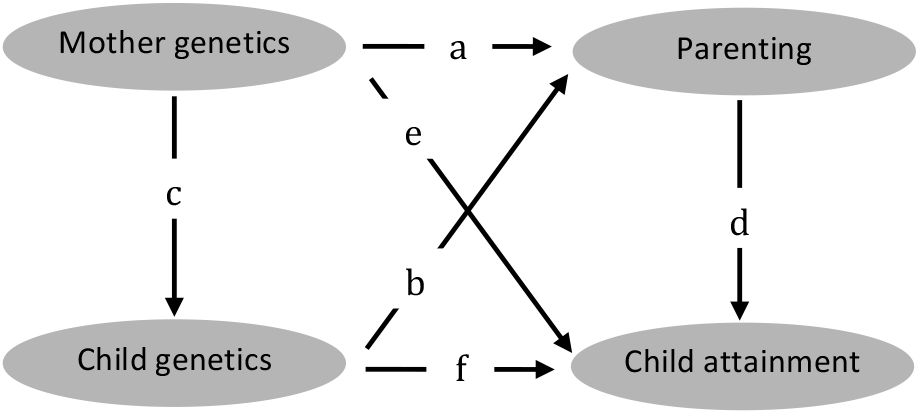
How do mothers’ and children’s education-associated genetics influence parenting and child attainment? Testing gene environment correlation, genetic confounding and genetic nurture.

Second, we tested for the presence of genetic confounding. We did this by testing whether associations between parenting and children’s educational attainment (Figure 1, Path d) reduce when controlling for children’s education polygenic scores. Genetic confounding as measured using the education polygenic score is possible (a) if mothers’ polygenic scores influence their parenting (Figure 1, path a) and the same genetics are passed on to children (Figure 1, path c) in whom they influence educational attainment (Figure 1, path f), or (b) if children’s polygenic scores both evoke the parenting they receive (Figure 1, path b) and also influence their educational attainment (Figure 1, path f). We therefore controlled for children’s polygenic scores to (a) control for education-associated genetics that influence parenting in the parent generation and that are passed on to children and (b) control for education-associated genetics in the child generation that evoke parenting (we did not additionally control for mothers’ education polygenic scores because confounding from mothers’ genetics can only arise if these genetics are passed on to children). Controlling for children’s polygenic scores does not entirely rule out genetic confounding, because the education polygenic score measures only a portion of all genetic influences on education. However, a finding that the association between parenting and children’s education reduces after controlling for children’s education polygenic score would support the hypothesis of genetic confounding, i.e. that parenting and children’s educational attainment are partly influenced by the same underlying genetic disposition.

Third, we tested for the presence of genetic nurture. We did this by testing whether mothers’ polygenic scores predicted their children’s educational attainment (Figure 1, path e). We tested this association controlling for children’s own polygenic scores, because mothers’ polygenic scores may predict children’s attainment simply due to mothers passing on genes to their children (Figure 1, path c*f). We previously reported in the E-Risk cohort that mothers’ polygenic scores predicted their children’s attainment over and above children’s own polygenic scores (Belsky et al., 2018). Here we directly tested the hypothesis that the parenting provided by mothers could explain this link between mothers’ polygenic scores and their children’s educational attainment (Figure 1, paths a*d). A finding that parenting explains the association would indicate that parental genetics affect children’s attainment independently of genetic transmission, via creating environments that influence children’s educational outcomes.

In summary, the goal of this article is to integrate new genetic discoveries into developmental psychology in order to test how genetics of both mothers and children influence the socialization context and children’s attainments.

## METHODS

### Participants

Participants were members of the Environmental Risk (E-Risk) Longitudinal Twin Study, which tracks the development of a birth cohort of 2,232 British children (Moffitt & E-Risk Study Team, 2002). Briefly, the E-Risk sample was constructed in 1999-2000, when 1,116 families (93% of those eligible) with same-sex 5-year-old twins participated in home-visit assessments. This sample comprised 56% monozygotic (MZ) and 44% dizygotic (DZ) twin pairs; sex was evenly distributed within zygosity (49% male). The study sample represents the full range of socioeconomic conditions in Great Britain, as reflected in the families’ distribution on a neighborhood-level socioeconomic index (ACORN [A Classification of Residential Neighborhoods], developed by CACI, Inc., for commercial use) (Odgers, Caspi, Russell, et al., 2012; Odgers, Caspi, Bates, Sampson, & Moffitt, 2012): 25.6% of E-Risk families live in “wealthy achiever” neighborhoods, compared with 25.3% nationwide; 5.3% compared with 11.6% in “urban prosperity” neighborhoods; 29.6% compared with 26.9% in “comfortably off” neighborhoods; 13.4% compared with 13.9% in “moderate means” neighborhoods; and 26.1% compared with 20.7% in “hard-pressed” neighborhoods. “Urban prosperity” families are underrepresented in E-Risk because such households are often childless.

Home visits were subsequently conducted when the children were aged 7 (98% participation), 10 (96%), 12 (96%), and at 18 years (93%). At age 18, 2,066 participants were assessed, each twin by a different interviewer. There were no differences between those who did and did not take part at age 18 in terms of socioeconomic status (SES) assessed when the cohort was initially defined (χ2=0.86, p=0.65), age-5 IQ scores (t=0.98, p=0.33), age-5 behavioral or emotional problems (t=0.40, p=0.69 and t=0.41, p=0.68, respectively). The Joint South London and Maudsley and the Institute of Psychiatry Research Ethics Committee approved each phase of the study. Parents gave informed consent and twins gave assent between 5-12 years and then informed consent at age 18.

### Parenting

We measured aspects of parenting that have previously been shown to predict children’s educational attainment: cognitive stimulation; warmth and sensitivity, household chaos (reverse-coded to indicate low household chaos), and safety and tidiness of the family home (Table 1). These aspects of parenting were assessed during home visits conducted at 4 time periods (when the children were aged 5, 7, 10, and 12 years of age) and drew on reports averaged across multiple informants - mothers, children and interview staff – to obtain comprehensive descriptions of the parenting children experienced during the first 12 years of their lives (Table 1). Parenting measures were positively correlated with each other (mean correlation r=.60, range .45-.70, all statistically significant at p<.01). Children’s educational attainment Children’s educational attainment was assessed in the age-18 interview, when children were asked to report their highest educational achievement. Educational attainment was classed following the Qualification and Credit Framework (QCF), a credit-based system used in the UK to assign educational qualifications to a set of ranked levels (http://www.accreditedqualifications.org.uk/qualifications-and-credit-framework-qcf.html). 18-year olds were classed as level 0 if they had no educational qualifications (3.4%); as level 1 if they scored a grade of D-G on their General Certificate of Secondary Education (GCSE) (18.5%); as level 2 if they scored a grade of A*-C (29.3%); and as level 3 if they had achieved or were currently working towards university entrance level qualifications (or equivalent) (48.9%).

**Table 1.** Description of parenting measures.

| Measure | Age | Informant | Format | Example items/statements |
| --- | --- | --- | --- | --- |
| **Cognitive stimulation** |  |  |  |  |
| Activities with mother | 5 | Mother | 12 items with 'yes'/'no' responses, reliability<sup>1</sup> $\alpha=.59$ | 'Have you and the children.. visited a zoo?' '...been to a park?' |
| Child stimulation | 7, 10, 12 | Interviewer | 6 items from the HOME<sup>2</sup>, with 'yes'/'a little'/'no' responses, mean reliability $\alpha=.77$ | 'Do the children have toys and puzzles?' 'Do the children have books?' |
| **Warm, sensitive parenting** |  |  |  |  |
| Parental warmth | 5, 10 | Mother | 5-minute speech sample<sup>3</sup>, interrater agreement $\kappa=.90$ | "She is a delight, she is so happy, I love taking her out, she is my ray of sunshine" |
| Parental dissatisfaction | 5, 10 | Mother | 5-minute speech sample<sup>3</sup>, interrater agreement $\kappa=.84$ | "I wish I had never had her . . . she's a cow, I hate her." |
| Positive parenting | 7, 10 | Interviewer | 5 items from the HOME<sup>2</sup>, with 'yes'/'a little'/'no' responses, mean reliability $\alpha=.82$ | 'Is the parent affectionate to the child?' 'Does the parent display warmth toward the child?' |
| Negative parenting | 7, 10 | Interviewer | 7 items from the HOME<sup>2</sup>, with 'yes'/'a little'/'no' responses, mean reliability $\alpha=.77$ | 'Is the parenting of the child overly strict?' 'Is the parent controlling towards the child?' |
| **Household chaos (reverse-coded)** |  |  |  |  |
| Interviewer report | 7, 10, 12 | Interviewer | 3 items from the HOME<sup>2</sup>, with 'yes'/'a little'/'no' responses, reliability $\alpha=.56$ | "Is the house chaotic or overly noisy?" |
| Mother and child report | 12 | Mother, child | 12 items with 'not'/'somewhat'/'very often or often true' responses, mean reliability $\alpha=.77$ | "You can hardly hear yourself think in our home" "We are always losing things at home" |
| **Safe, tidy home** |  |  |  |  |
| | 5, 7, 10, 12 | Interviewer | 2-4 items (depending on age) with 'yes'/'a little'/'no' responses, mean reliability $\alpha=.79$ | 'Did the home or flat appear safe?' 'Are visible rooms of the house clean?' |

<sup>1</sup> Internal consistency reliability (Cronbach's alpha). <sup>2</sup> Home Observation for Measurement of the Environment (HOME) (Caldwell & Bradley, 1984). <sup>3</sup> Using procedures adapted from the Five Minute Speech Sample method (Magaña et al., 1986) as described previously (Caspi et al., 2004).

### Genotyping and imputation

We used Illumina HumanOmni Express 24 BeadChip arrays (Versions 1.1 and 1.2; Illumina, Hayward, CA) to assay common single-nucleotide polymorphism (SNP) variation in the genomes of E-Risk participants and their mothers. We imputed additional SNPs using the IMPUTE2 software (Version 2.3.1, https://mathgen.stats.ox.ac.uk/impute/impute_v2.html; Howie, Donnelly, & Marchini, 2009) and the 1000 Genomes Phase 3 reference panel (Abecasis et al., 2012). Imputation was conducted on SNPs appearing in dbSNP (Version 140; http://www.ncbi.nlm.nih.gov/SNP/; Sherry et al., 2001) that were “called” in more than 98% of the samples. Invariant SNPs were excluded. The E-Risk cohort contains monozygotic twins, who are genetically identical; we therefore empirically measured genotypes of one randomly-selected twin per pair and assigned these data to their monozygotic co-twin. Prephasing and imputation were conducted using a 50-million-base-pair sliding window. The resulting genotype databases included genotyped SNPs and SNPs imputed with 90% probability of a specific genotype among European-descent members of the E-Risk cohort. We analyzed SNPs in Hardy-Weinberg equilibrium (*p* > .01). We restricted our analyses to European-descent study participants because allele frequencies, linkage disequilibrium patterns, and environmental moderators of associations may vary across populations (Martin et al., 2017). Of the N=1,116 E-Risk families, there were n=860 families for whom family members’ genetic data could be analyzed, based on mothers and at least one child having genetic data. In families with versus without genetic data there were no differences in parenting, but children’s educational attainment tended to be lower among those for whom genetic data was analyzed (p=.05).

### Polygenic scoring

Polygenic scoring was conducted following the method described by Dudbridge (Dudbridge, 2013) using PRSice (Euesden, Lewis, & O’Reilly, 2015). Briefly, SNPs reported in the most recent GWAS results released by the Social Science Genetic Association Consortium (Lee et al., 2018) were matched with SNPs in the E-Risk database. For each SNP, the count of education-associated alleles was weighted according to the effect estimated in the GWAS. Weighted counts were averaged across SNPs to compute polygenic scores. We used all matched SNPs to compute polygenic scores irrespective of nominal significance for their association with educational attainment and linkage disequilibrium between SNPs. To control for possible population stratification, we conducted a principal components analysis of our genome-wide SNP database using PLINK v1.9 (Chang et al., 2015). We residualized polygenic scores for the first ten principal components estimated from the genome-wide SNP data. The residualized score was normally distributed and standardized to M=0, SD=1.

### Statistical analysis

We used structural equation models for dyads with indistinguishable members (Kenny, Kashy, & Cook, 2006) to test gene-environment correlation, genetic confounding and genetic nurture. In these models, analyses are conducted at the family level while constraining means and corresponding paths for twins to be equal. To test gene-environment correlation, we fitted a model predicting parenting from mothers’ and children’s education polygenic scores, first each separately, then all together in the same model. To test genetic confounding, we fitted a model predicting children’s educational attainment from parenting, and tested whether associations between parenting and educational attainment reduced when accounting for children’s polygenic scores. To test genetic nurture, we fitted a model predicting children’s educational attainment from mothers’ education polygenic scores, and added children’s polygenic scores to this model to test effects of mothers’ polygenic scores over and above children’s own scores. We then added the parenting variables to this genetic-nurture model as mediators. Each parenting variable was initially tested separately, and then all significant mediators were entered together into the same model. We adjusted for children’s sex in all analyses. All measures were standardized to M=0, SD=1. All analyses were conducted using Mplus version 8.2 (Muthén & Muthén, 1998-2017).

## RESULTS

### Both nature and nurture predict children’s educational attainment

As expected, children’s education polygenic scores predicted their educational attainment: children with higher polygenic scores completed higher levels of education (β=.24 [95%CI .19, .30], p<.01). Also as expected, parenting predicted children’s educational attainment: children exposed to greater cognitive stimulation, more warm, sensitive parenting, less household chaos and a safer, tidier home environment went on to complete more education (estimates ranged from β=.33 for safe, tidy home environment to β=.52 for cognitive stimulation; Figure 2).

**Figure 2.**
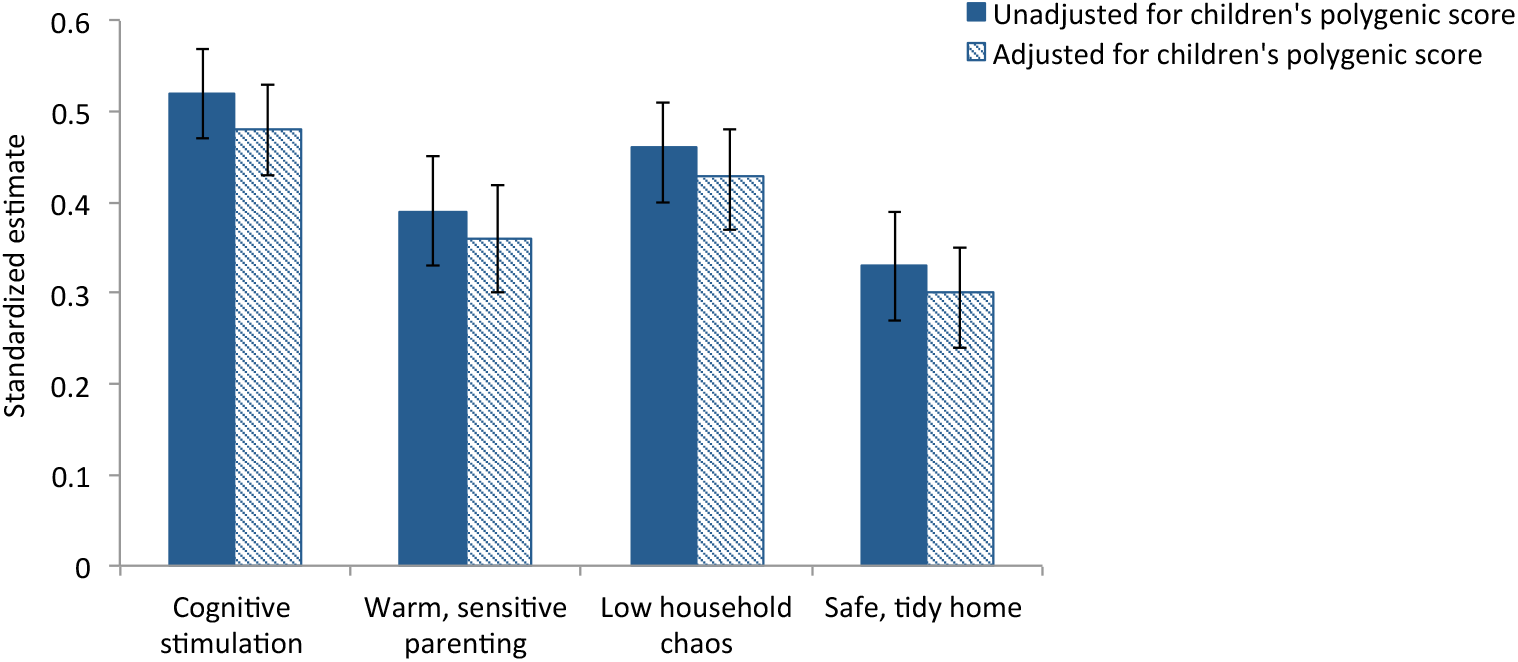
Genetic confounding: Controlling for children’s polygenic scores slightly reduces the prediction from parenting to children’s educational attainment.

### Testing gene-environment correlation: Do mothers’ and children’s education polygenic scores predict parenting?

Our results provided evidence for gene-environment correlation. Mothers with higher education polygenic scores provided greater cognitive stimulation and more warm, sensitive parenting, and raised their children in less chaotic and safer, tidier homes (estimates ranged from β=.14 for warm, sensitive parenting to β=.25 for cognitive stimulation; Figure 3). Children with higher polygenic scores also received more cognitive stimulation and more warm, sensitive parenting, and were raised in less chaotic and safer, tidier homes (estimates ranged from β=.12 for safe, tidy home to β=.21 for cognitive stimulation; Figure 3). As would be expected, mothers’ and children’s education polygenic scores were correlated (β=.51 [95%CI .46, .56], p<.01), due to mothers passing on genes to their children. We therefore included mothers’ and children’s education polygenic scores in the same models in order to predict each of the parenting behaviours. In these models, mothers’ education polygenic scores predicted all aspects of parenting independently of their children’s polygenic scores, indicating active gene-environment correlations between mothers’ genetics and parenting (Figure 3). In addition, children’s polygenic scores predicted cognitive stimulation and warm, sensitive parenting independently of their mothers’ polygenic scores, indicating evocative gene-environment correlations between children’s genetics and these aspects of parenting. Children’s polygenic scores did not predict household chaos or the safety and tidiness of the home independently of their mothers’ polygenic scores, indicating that these aspects of parenting are shaped more by mothers’ than children’s education-associated genetics (Figure 3).

**Figure 3.**
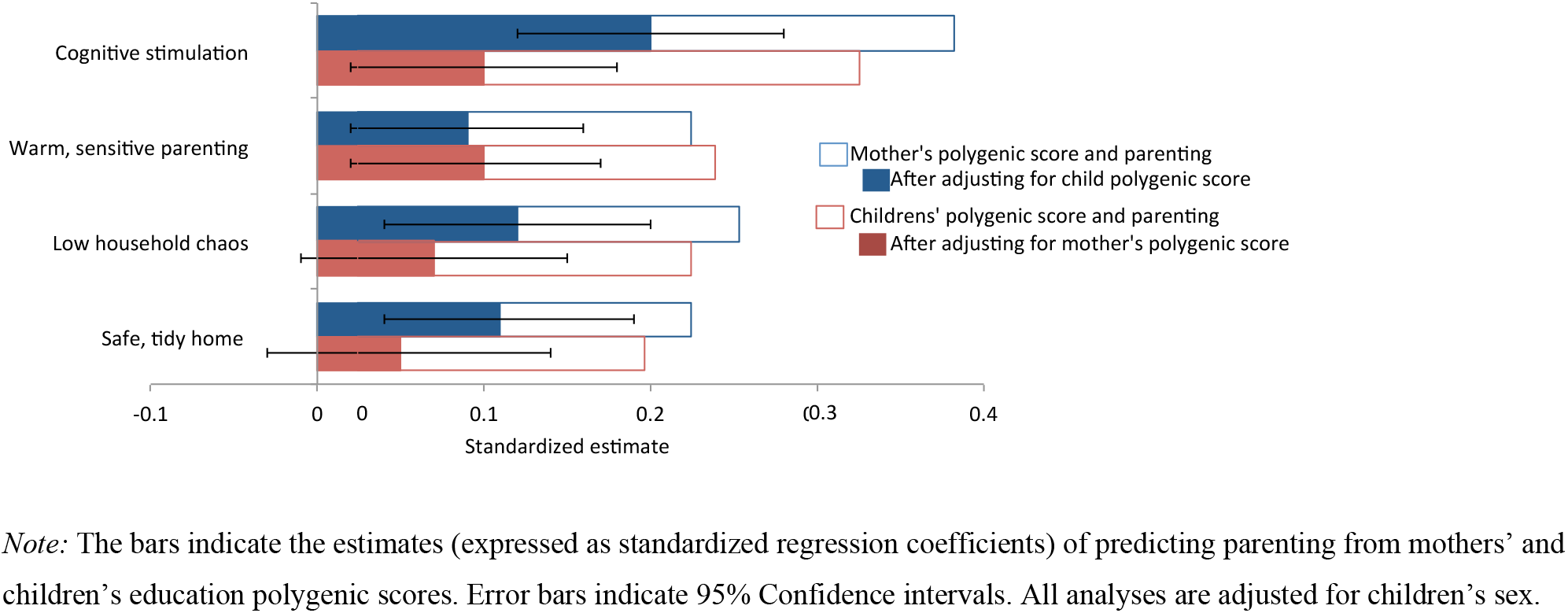
Gene-environment correlation: Mothers’ and children’s education polygenic scores predict parenting.

### Testing genetic confounding: Do genetic influences confound associations between parenting and children’s education?

Our results provided modest evidence for genetic confounding. Without controlling for genetics, children exposed to greater cognitive stimulation, more warm, sensitive parenting, less household chaos and a safer, tidier home environment went on to complete more education (estimates ranged from β=.33 for safe, tidy home environment to β=.52 for cognitive stimulation; Figure 2). Controlling for genetics led to a significant reduction of these associations, by approximately 7%, but the attenuations were small and parenting continued to be a statistically significant predictor of educational attainment. These findings indicate that genetic influences, as captured by the education polygenic score, account for only a small part of the reason for why parenting predicts children’s educational attainment.

### Testing genetic nurture: How do maternal genetics and parenting combine to influence children’s education?

Our findings provided evidence for genetic nurture. We first tested whether mothers’ education polygenic scores predicted their children’s educational attainment; this was the case (β=.23 [95%CI .17, .29], p<.01). The association was not simply due to mothers passing on education-associated genetics to their children; mothers’ education polygenic scores predicted their children’s educational attainment over and above children’s own polygenic scores (β=.12 [95%CI .05, .19], p<.01). This finding suggests the hypothesis that mothers’ education-associated genetics shape family environments that affect children’s attainments independently of mother-child genetic transmission. We tested this hypothesis by adding measures of parenting to the analysis model. Of the four parenting measures, three (cognitive stimulation, household chaos, and a safe, tidy home) emerged as statistically significant mediators (Table 2). Cognitive stimulation on its own accounted for approximately 75% of the association between maternal genetics and children’s educational attainment. Low household chaos and a safe, tidy home each mediated approximately 42% and 25% of the association, respectively, but in a model containing cognitive stimulation, only low household chaos accounted for a small portion of additional covariance beyond cognitive stimulation. These findings indicate that mothers’ education polygenic scores influence their children’s attainment via mothers’ parenting, particularly the extent of cognitive stimulation mothers provided to their children.

**Table 2.** Genetic nurture: Parenting mediates associations between mothers’ education polygenic scores and their children’s educational attainment independently of children’s polygenic scores.

|  | Parenting |  |  |  |  |
| --- | --- | --- | --- | --- | --- |
|  | Cognitive stimulation | Warm, sensitive parenting | Low household chaos | Safe, tidy home | All significant<sup>2</sup> mediators together |
|  | Estimate (95% CI)<sup>1</sup> | Estimate (95% CI) | Estimate (95% CI) | Estimate (95% CI) | Estimate (95% CI) |
| Total effect | **.12 (.05, .19)** | **.12 (.05, .19)** | **.12 (.05, .19)** | **.12 (.05, .19)** | **.12 (.05, .19)** |
| Direct effect | .03 (-.03, .09) | **.10 (.03, .16)** | **.07 (.01, .14)** | **.09 (.03, .16)** | .03 (-.03, .09) |
| Total indirect effect | **.09 (.05, .13)** | .02 (.00, .04) | **.05 (.01, .08)** | **.03 (.01, .06)** | **.09 (.05, .13)** |
| % Mediation | **75%** | 17% | **42%** | **25%** | **75%** |

<sup>1</sup> Indicates standardized estimates of total, direct and indirect effects in mediation models. CI=confidence intervals. Bolded estimates indicate statistically significant effects.
<sup>2</sup> Model includes cognitive stimulation, low household chaos and safe, tidy home.

## DISCUSSION

The investments parents make to raise their offspring are thought to be a major contributor to children’s educational success, making parental investment a cornerstone of psychological, sociological, and economic models that seek to explain how educational inequalities are created and perpetuated (Cheng, Johnson, & Goodman, 2016; Feinstein, Duckworth, & Sabates, 2004; Kalil, 2015). However, findings from behavior-genetic studies have challenged causal interpretations of parental influence by showing genetic influences on parenting; a gene-environment correlation. Here we tested implications of gene-environment correlations for parental investment in children’s educational attainment using a novel design: In a prospective-longitudinal study, we collected genotype data from both mothers and children and matched these genetic data with home-visit measures of parenting behavior. We report three main findings.

First, we found evidence for gene-environment correlations. Both mothers’ and children’s education-associated genetics, summarized in genome-wide polygenic scores, predicted the kind of parenting that is known to be linked with children’s later educational success. By collecting genetic data from both mothers and their offspring we were able to show that different forms of gene-environment correlations operate in the same family, at the same time. Active- and evocative gene-environment correlations were both implicated in the cognitive stimulation and the warm, sensitive parenting that children experienced. In addition, active gene-environment correlations were also implicated in the kinds of households (chaotic, safe, tidy) in which children grew up. Second, we found evidence for slight genetic confounding. The estimated effects of mothers’ parenting on children’s educational attainment were significantly reduced after accounting for education-associated genetics, consistent with a view of genes as confounding part of the link between parenting and child attainment. However, the magnitude of the confounding was small. Third, we found evidence for genetic nurture. Parenting behavior -- particularly mothers’ cognitive stimulation of their children -- explained why mothers’ genetics influenced their children’s educational attainment (over and above genetic transmission from mother to child). This finding extends recent reports of associations between parental genetics and children’s educational attainment (Bates et al., 2018; Belsky et al., 2018; Kong et al., 2018; Liu, 2018) by showing, for the first time, that parents’ education-associated genetics actively shape features of the family environment that influence the next generation’s educational success.

Our findings need to be interpreted in light of several limitations. First, our approach to estimating genetic nurture relies on the assumption that mothers’ and children’s polygenic scores are measured with identical error (Belsky et al., 2018). To the extent that this assumption is violated, our estimates of genetic nurture could be upwardly or downwardly biased, depending on whether error is greater in mothers’ versus children’s polygenic scores. However, the assumption is probably defensible, because mothers’ and children’s polygenic scores are identical measurements, i.e. sums of the same genotypes transformed using the same weights. Second, although the education polygenic score that we used is based on the largest-ever social-science GWAS, a limitation of this GWAS is that it still reflects only a portion of all genetic influences on educational attainment (approximately one third) (Lee et al., 2018). To the extent that the polygenic score is an underestimate of the total genetic influence on educational attainment, our estimates of gene-environment correlations, genetic confounding, and possibly genetic nurture are likely to be underestimates of the true effects. At this point, our findings provide ‘proof-of-principle’ of these processes, and the implications they raise can continue to be tested as refined polygenic scores become available. Third, we tested genetic confounding and genetic nurture only for children’s educational attainment, not for other child outcomes. We focused on educational attainment because it is a central determinant of future health, wealth and wellbeing (Cutler & Lleras-Muney, 2010; Hout, 2012; Oreopoulos & Salvanes, 2011), and because the polygenic score for educational attainment is based on the largest GWAS of a social-behavior phenotype (Lee et al., 2018). To the extent that other developmental outcomes are both genetically-influenced and associated with parenting, processes of genetic confounding and genetic nurture will likely be present as well. As increasingly-larger GWAS are conducted for more developmental outcomes, the same design we present here can be used to test genetic confounding and genetic nurture for these outcomes. Fourth, we did not have genetic data from fathers, which means that we were unable to control for fathers’ education polygenic scores when estimating associations between mothers’ and children’s education polygenic scores and parenting. To the extent that fathers’ genes are correlated with parenting, the associations we observed in our study may therefore partly reflect effects of fathers’ genetics, because fathers’ and children’s genes are correlated (due to genetic inheritance) and because mothers and fathers’ genetics may be correlated (due to assortative mating, i.e. the tendency to select partners with characteristics similar to one’s own). Against this background, we conclude by discussing the implications of our findings about (a) gene-environment correlations, (b) genetic confounding, and (c) genetic nurture for a more thorough understanding of the developmental processes that shape children’s attainment.

Our findings of gene-environment correlation replicate and extend our prior work on genetic associations with parenting (Wertz et al., submitted). We replicated findings from a previous analysis in a New Zealand cohort, in which we showed that parents’ education-associated genetics shape the warm, sensitive, stimulating parenting they provide to their children (Wertz et al., submitted). Here we report the same pattern of results in an independent cohort of British mothers, indicating that genetic correlations with parenting are robust against differences in context and measurements of parenting. We extend this prior work by incorporating children’s polygenic scores in our analyses, finding that children’s education-associated genetics shape the parenting they receive. Together with other recent studies (Dobewall et al., 2018; Krapohl et al., 2017; Selzam et al., 2018), these findings provide molecular-genetic evidence for a bidirectional model of parent-child relations, in which parenting is partly a response to children’s characteristics (Bell, 1968; Crouter & Booth, 2003; Pardini, 2008; Sameroff, 2010).

Findings of gene-environment correlations with parenting imply that family environments children experience while growing up are partly a function of their own and their parents’ genetics. For example, we found that children of parents who carried a high number of education-associated variants were exposed to greater cognitive stimulation in the home compared to children of parents who carried fewer of these variants. Because parents and children share genes, family environments shaped by parents’ genes will tend to match and reinforce children’s genetic dispositions (Knafo & Jaffee, 2013; Scarr & McCartney, 1983; Tucker-Drob & Harden, 2012). Such a match can positively influence children’s development; for example, when a child with a high education polygenic score is born into a family that provides cognitive stimulation. However, the same match also implies that a child genetically at-risk for poor educational outcomes will tend not to experience exactly the kind of stimulating and supportive parenting that could make a difference for his or her attainment. Thus, for better and for worse, correlations between genes and environments reduce the availability of experiences that can alter individuals’ developmental trajectories. This also applies to the reproduction of educational success across generations. To the extent that educational outcomes are influenced by genetics, genes will tend to be a force for intergenerational stability in educational attainment, both via direct genetic transmission and via indirect effects of genes on caregiving environments that shape future generations’ behaviors. This tendency means that is it important to improve children’s access to interventions that may be able to break reinforcing links between genes and environments, such as high-quality early learning programs (Heckman, 2006).

Given how much attention critics of parenting effects devote to the possibility of genetic confounding (Harris, 1998; Rowe, 1993; Sherlock & Zietsch, 2018), it may seem surprising that our estimates of genetic confounding were so small. There are two possible explanations for this finding: either genetics do little to confound associations between parenting and children’s educational attainment, or we have underestimated the true magnitude of genetic confounding. The observation that polygenic-score associations with educational attainment are substantially lower than heritability estimates of educational attainment (Branigan, McCallum, & Freese, 2013) suggests that our findings most likely underestimate genetic confounding. Currently, even the best and biggest efforts to capture the genetic variants influencing educational attainment are still missing a substantial part of its heritability (Manolio et al., 2009; National Human Genome Research Institute, 2018). Until more of this ‘missing heritability’ can be accounted for at the molecular genetic level, the safest way to rule out genetic confounding is to continue to use family-based designs such as the discordant-twin design (McGue, Osler, & Christensen, 2010; Vitaro, Brendgen, & Arseneault, 2009), parent-child adoption design (Leve et al., 2013) or children-of-twin design (D’Onofrio et al., 2003), that can estimate associations between parenting and children’s educational attainment free from genetic influences shared between parents and children (Turkheimer & Harden, 2014).

Debates about parental influences on children’s development tend to contrast the effects of parents’ genes -- assumed to influence children via genetic transmission -- with the effects of parenting -- assumed to influence children via environmental ways. Our finding of genetic nurture draws a more nuanced picture, showing that mothers’ genes affect children’s attainment over and above genetic transmission, via parenting. This finding has three implications. First, over and above a person’s own genetics, their development will be shaped by the genomes of significant others. We demonstrate this here for effects of a parents’ genetics on children’s outcomes, but this observation likely extends beyond parents to everyone who creates environments inhabited by people – family members; individuals residing outside the family context, such as peers and partners (Conley et al., 2016; Domingue et al., 2018); even people a child may be exposed to only indirectly, such as the grandparents who raised a child’s parents (Hällsten & Pfeffer, 2017; Kong et al., 2018; Liu, 2018). The existence of a ‘social genome’ broadens the scope of the study of genetics, from an individual’s genes and their effects on an individual’s phenotype, to the genome of an individual’s social context (Domingue & Belsky, 2017). Second, much has been written about the need to integrate genetics into parenting research and socialization theory, but there is also a need to integrate environments into how to think about and collect genetic data. Correlations between genes and environments are a challenge not only for socialization research, but also for genetics research: Although DNA sequence cannot be modified by the environment, our findings show that environments still pose a threat to causal inference, because associations between a person’s DNA and developmental outcomes may partly reflect effects of environments created through genes of other individuals (Kong et al., 2018)). As much as genetic confounding needs to be considered when estimating environmental effects, ‘environmental confounding’ needs to be taken into account when estimating genetic effects (Krapohl et al., 2017; Young et al., 2018). Third, our findings show that environments are part of the pathway from genotype to phenotype (Kandler & Zapko-Willmes, 2017; Scarr & McCartney, 1983). Specifically, we found that genetic influences on children’s educational attainment partly manifested through parenting; an environmentally mediated genetic effect. Combining genetic data with measures of individuals’ social environments is key to tracing how genetics affect life outcomes. By joining forces in this way, genetics and socialization researchers will be able to strengthen causal estimates and obtain a more complete understanding of the processes shaping children’s attainments.

## ACKNOWLEDGEMENTS

The Environmental Risk (E-Risk) Longitudinal Twin Study is funded by U.K. Medical Research Council (UKMRC grant G1002190). Additional support was provided by the U.S. National Institute of Child Health and Development (NICHD) grant HD077482 and by the Jacobs Foundation. L. Arseneault is the Mental Health Leadership Fellow for the UK Economic and Social Research Council (ESRC). J. Agnew-Blais is a UK Medical Research Council Skills Development Fellow. D.W. Belsky is supported by an Early-Career Research Fellowships from the Jacobs Foundation. L.S. Richmond-Rakerd is a postdoctoral fellow with the Carolina Consortium on Human Development and the Center for Developmental Science at the University of North Carolina-Chapel Hill. We are grateful to the study mothers and fathers, the twins and the twins’ teachers for their participation. Our thanks to members of the E-Risk team for their dedication, hard work and insights.

